# Impact of global warming on insects: are tropical species more vulnerable than temperate species?

**DOI:** 10.1101/728352

**Authors:** Frank Johansson, Germán Orizaola, Viktor Nilsson-Örtman

## Abstract

The magnitude and ecological impact of climate change varies with latitude. Several recent models have shown that tropical ectotherms face the greatest risk from warming because they currently experience temperatures much closer to their physiological optimum than temperate taxa. Even a small increase in temperature may thus result in steep fitness declines in tropical species but increased fitness in temperate species. This prediction, however, is based on a model that does not account for latitudinal differences in activity periods. Temperate species in particular may often experience considerably higher temperatures than expected during the active season. Here, we integrate data on insect warming tolerance and temperature-dependent development to re-evaluate latitudinal trends in thermal safety margins after accounting for latitudinal trends in insect seasonal activity. Our analyses suggest that temperate and tropical species differ far less in thermal safety margins than commonly assumed, and thus face a similar risk from warming.

## Introduction

Global warming is expected to have a severe impact on the earth’s biodiversity and ecosystems (Urban 2015). In fact, numerous studies have documented pronounced shifts in the phenology, physiology and distribution of plant and animal species associated with recent changes in climatic conditions (Parmesan 2006, 2007, Dillon et al. 2010, Thackeray et al. 2016, Pecl et al. 2017, Lister and Garcia 2018). As a consequence, there is an urgent need to develop methods that can predict the impact of future climate change on populations and species at a global scale. Ideally, these methods should be able to identify broad geographic or taxonomic trends in the susceptibility of organisms to global warming, thereby helping to guide efforts to alleviate the consequences of global warming more effectively (Root et al. 2003, Parmesan 2007). While considerable progress has been made toward developing such a framework (Thuiller 2004, Botkin et al. 2007, Deutsch et al. 2008, Sinclair et al. 2016), many challenges remain.

In this emerging framework, thermal performance curves (TPCs) have become a key component (Figure 1). TPCs describe how temperature affects an organisms’ fitness or key contributing functions, such as locomotion, growth and reproduction (Huey and Stevenson 1979, Sinclair et al. 2016). TPCs are especially relevant for ectotherms – the most diverse and widespread group of terrestrial animals – because ambient temperature has a direct and profound impact on nearly all aspects of ectotherm performance. TPCs for ectotherm performance and fitness typically show a gradual increase in performance from a critical thermal minimum (T_min_) where performance or fitness is zero, until reaching an optimal temperature (T_opt_) where performance is highest, before decreasing rapidly towards a critical thermal maximum (T_max_) (Figure 1).

**Figure 1.**
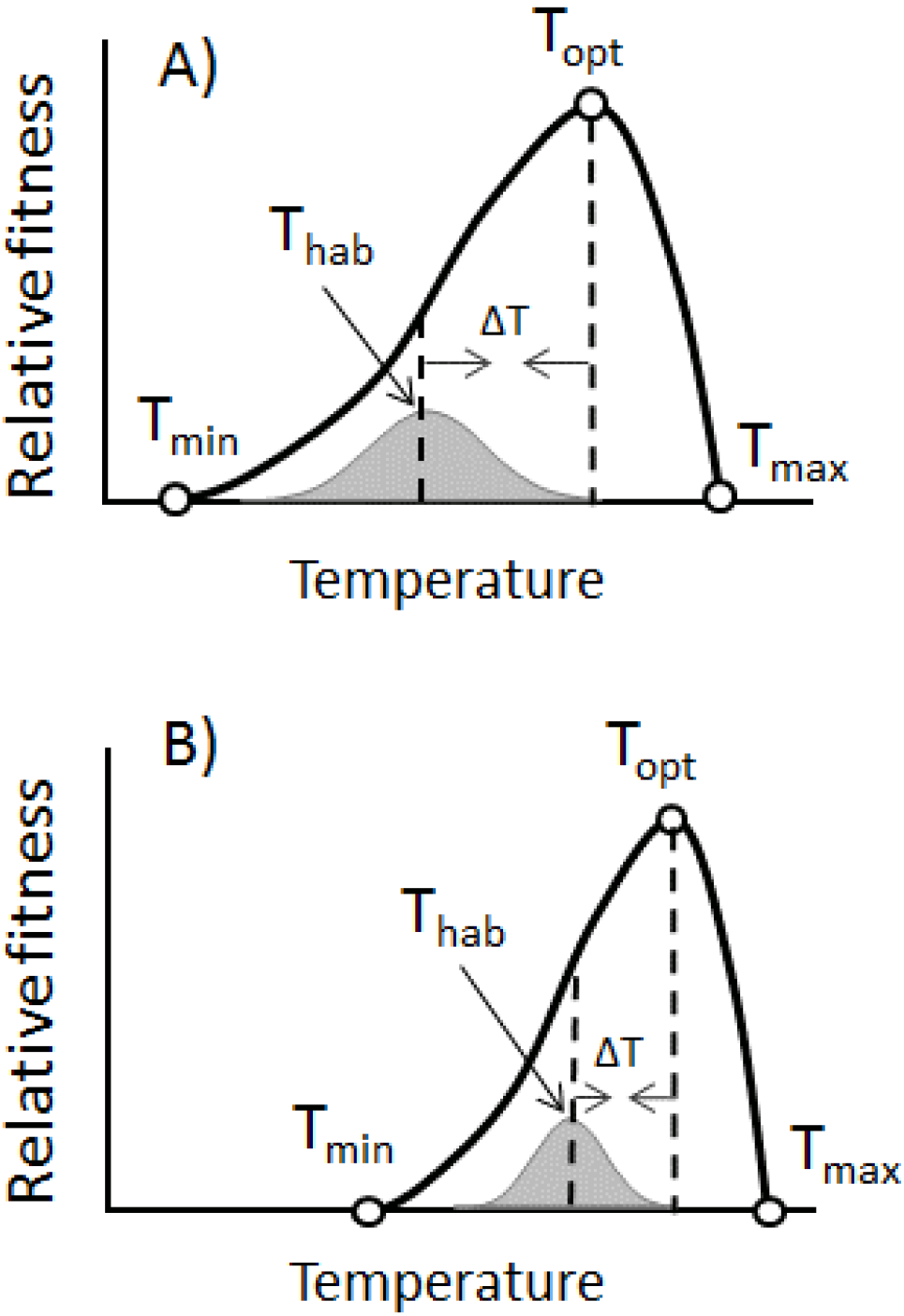
Thermal performance curves of ectotherms, here depicted as relative fitness. A and B represent a temperate and a tropical species, respectively. T_min_ and T_max_ represent the minimum and maximum temperature at which organisms can perform, and T_opt_ is the optimal temperature for performance. The grey curve beneath the thermal performance curve is the temperature that the organism is exposed to during an average year and the average is depicted as T_hab_. ΔT is the distance between T_hab_ and T_opt_. With global warming this distance decreases and might even be shifted to the right of T_opt_ or T_max_ (Deutch et al 2008). Tropical species are assumed to be temperature specialists (B), because they have narrower thermal performance curves and, therefore, are predicted to be more sensitive to global warming (Deutsch *et al.* 2008).

Recently, several studies have synthesized and analyzed ectotherm TPC data on a global scale. Most strikingly, these studies have brought widespread attention to the fact that tropical species appear to be more vulnerable to global warming than temperate species. For example, in a seminal paper, Deutsch et al. (Deutsch et al. 2008) used TPCs for insect fitness to estimate two measures of an organism’s susceptibility to warming: the amount of warming that an organism can tolerate before it experiences a decline in fitness (thermal safety margin: TSM) or reach zero fitness (warming tolerance: WT). The results revealed that tropical species have considerably narrower thermal safety margins and warming tolerances than temperate species (Deutsch et al. 2008). This suggests that even a small increase in global mean temperatures will result in precipitous declines in the fitness and performance of tropical species – as they become pushed toward or beyond their thermal maxima – whereas temperate species will benefit from an increase in mean temperatures, as they currently experience temperatures well below their thermal optimum (Addo-Bediako et al. 2000, Compton et al. 2007, Sunday et al. 2011). Later studies, employing a similar methodology, have broadly supported these conclusions (Tewksbury et al. 2008, Huey et al. 2009, Dillon et al. 2010, Diamond et al. 2012, Duarte et al. 2012, Kingsolver et al. 2013).

However, this prediction, that tropical species are more vulnerable to warming than temperate taxa, rests on several important assumptions. Here, we examine one of these assumptions, namely that the duration of the active period of insects is similar across latitudes. This assumption arises from using annual climatological data when performing these analyses. In other words, in many studies, estimates of T_opt_ and T_max_ are compared with the mean habitat temperature across the year at each location. However, because most ectotherms become inactive during the parts of the year when conditions are unfavorable (Leather et al. 1995), most organisms will tend to experience warmer and less variable temperatures during the active season than expected based on annual means and variances. Deutsch et al. (Deutsch et al. 2008) was aware of this, and complemented their original analyses (based on annual mean temperatures) with an analysis were they calculated WTs and TSMs using the mean temperature during the three warmest months of the year at each location. Overall, this did not change their conclusions regarding latitudinal trends in the susceptibility to warming. However, in these complementary analyses, temperate and tropical organisms were assumed to have identical seasonal activity patterns (and thus become inactive during a major part of the year). This contrasts with the empirical evidence, which shows clear latitudinal trends in ectotherm activity patterns (Leather et al. 1995, Buckley et al. 2017). In particular, temperate insects display a much stronger association between insect activity and temperature, meaning that the mismatch between the annual climate and that experienced by individuals during the active season is likely to be greatest for insects from higher latitudes. Whether restricting the analysis to the three warmest months of the year (as in (Deutsch et al. 2008) or the six warmest months (as in (Kingsolver et al. 2013) re-analysis of the Deutsch et al. dataset) represents a realistic approximation of the thermal environment experienced by active insects across latitude remains unknown. Taking the biologically active period of insects and other ectotherms into consideration could thus be critical for generating more precise and biologically relevant predictions for the effects of global warming across latitudes. However, to our knowledge, no study has assessed the consequences of empirically-observed latitudinal differences in activity periods on the vulnerability to warming at a global scale.

Here, we re-examine the prediction that tropical species currently experience mean habitat temperatures that are much closer to their thermal optimum and maximum than temperate species and thus are at a greater risk from warming. To do this, we revisit the dataset used by Deutsch et al. (Deutsch et al. 2008) and test for differences between tropical and temperate species in the vulnerability to global warming, after accounting for empirically-based estimates of insect active periods across latitudes. Admittedly, our analyses – as well as those of Deutsch et al (2008) – ignore several additional factors that are increasingly known to be important for ectothermic responses to global warming, including thermoregulatory behavior (Buckley et al. 2015), capacity for thermal acclimation (Nilsson-Örtman and Johansson 2017), non-linear effects of temperature variance (Vasseur et al. 2014), response to the duration and intensity of extreme temperatures (Marshall and Sinclair 2015), shifts in phenology (Visser and Both 2005, Buckley et al. 2017), genetic (co)variances (van Heerwaarden et al. 2016) and interspecific interactions (Nilsson-Örtman et al. 2014). Nevertheless, by focusing specifically on the assumption that active periods are similar across latitudes, we explore the importance of accounting for insect activity periods when deriving predictions for the vulnerability of populations and species across latitude to future global warming.

We define the active period of each studied insect population as the months of the year at a given location when the average temperature falls above the lower thermal threshold for insect development. After establishing the active period at each location, we re-calculate warming tolerances (WT) and thermal safety margins (TSM) using both the annual mean habitat temperature (T_hab_) originally used by Deutsch et al. (Deutsch et al. 2008)) and a new metric described here; the annual mean habitat temperature during the active period T_habA._ When we account for differences in insect activity periods across latitudes, we predict that: 1) temperate species will experience temperatures much closer to their T_opt_ and T_max_ than previously assumed (Figure 1); 2) temperate and tropical species will thus have more similar warming tolerances and thermal safety margins than expected based on the mean annual temperature; and 3) temperate and tropical species will face a similar risk of experiencing fitness declines under future global warming scenarios.

**Figure 2.**
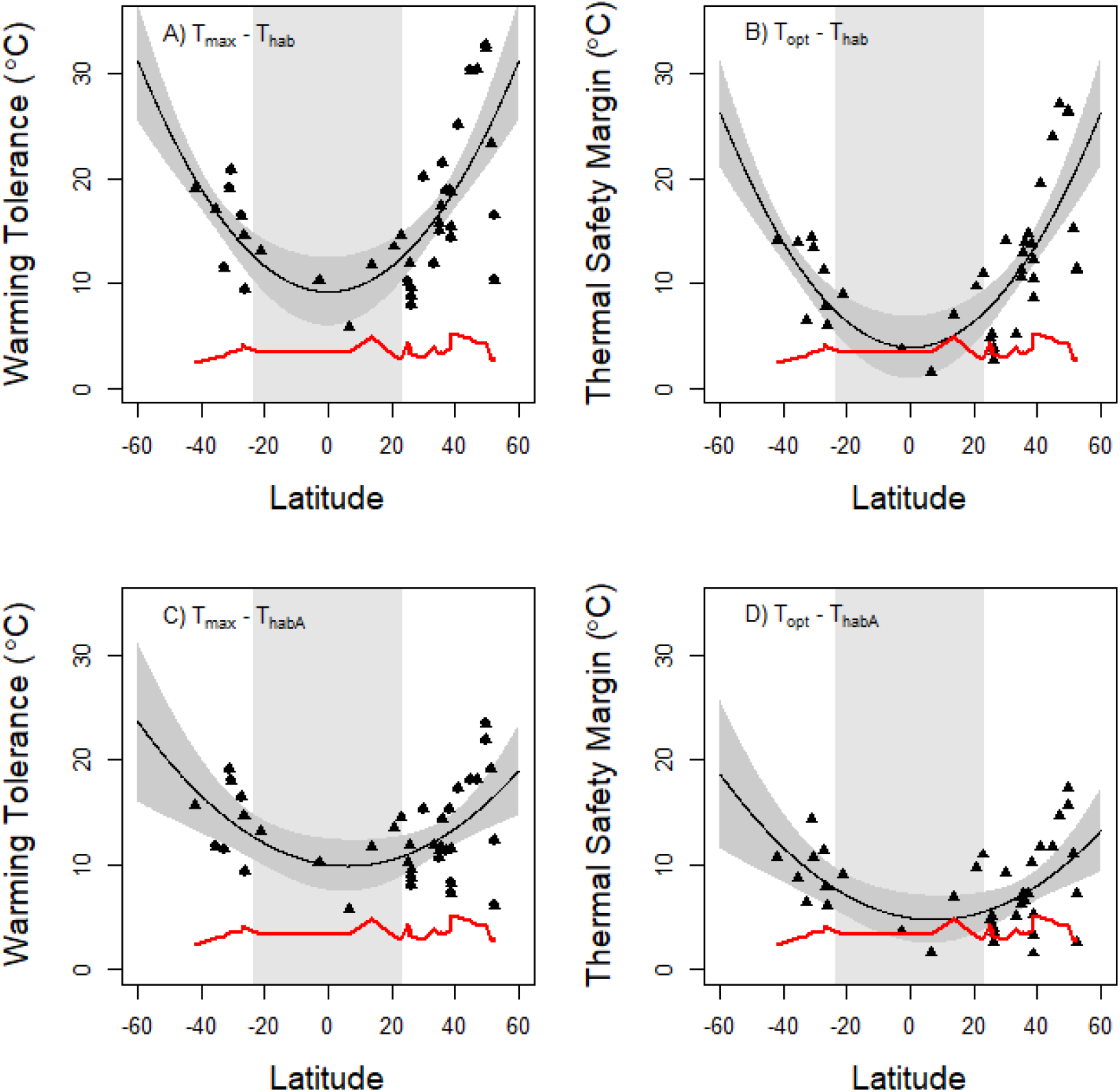
Warming tolerances (WT) and thermal safety margins (TSM) of insects as a function of latitude. In A and B, warming tolerances and thermal safety margins were calculated using on the mean annual temperature Thab as in several previous studies (i.e. WT = T_max_−T_hab_; TSM = T_opt_−T_hab_). In C and D, warming tolerances and thermal safety margins were calculated using the mean temperature during the months of the year when insects are developmentally active, T_habA_ (i.e. WT = T_max_−T_habA;_ TSM = T_opt_−T_habA_). The red line represents the predicted increase in mean temperature at each latitude by 2080. Species whose data points fall below this line in B and D are predicted to experience mean habitat temperature that exceed their T_opt_ in 2080, and thus decline in fitness. Note that species whose data points fall near the red line may also experience fitness declines under climate change due to thermal fluctuations that exceed T_opt_. Tropical areas (defined here as regions located between 23°N and 23°S latitude) are shaded in light gray. Predictions from a second-degree polynomial model for the latitudinal trend in WT and TSM are shown in black; 95% confidence intervals for the predictions are shown in dark gray.

## Material and methods

We combine two global eco-physiological datasets to re-evaluate the impact of global warming on insects across latitudes: the dataset on insect thermal tolerance previously analyzed by Deutsch et al. (Deutsch et al. 2008) and a dataset on temperature thresholds of insect development published by Dixon et al. (Dixon et al. 2009). Importantly for this study, the Deutsch et al. (Deutsch et al. 2008) dataset contain estimates of T_opt_ and T_max_ derived from TPCs for insect fitness (intrinsic population growth rate) from 38 insect species globally, and the Dixon et al. (Dixon et al. 2009) dataset contain estimates of the minimum developmental temperature for 66 species of insects. Briefly, our analysis followed a four-step procedure. First, we defined the lower thermal threshold for insect development (T_dmin_) using data from Dixon et al. (Dixon et al. 2009). Second, we defined the active period of each insect population in the Deutsch et al. (Deutsch et al. 2008) dataset as the months of the year when the mean habitat temperature fell above T_dmin_. Third, we calculated a novel metric, T_habA_, which we defined as the mean habitat temperature during the active period for each population in the Deutsch et al. (Deutsch et al. 2008) dataset. Finally, we re-calculated the warming tolerance WT as T_max_-T_habA_ and the thermal safety margin TSM as T_opt_-T_habA_ for each population and plotted these metrics against latitude. We describe the analysis in greater detail next.

### Minimum developmental temperature (T_dmin_)

Insects cannot develop below a certain temperature (Dixon et al. 2009). Therefore, the majority of insects at high latitudes are only active during the warmer months in a year (Leather et al. 1995) and enter diapause at some point near the end of the growth season (Tauber et al. 1986, Leather et al. 1995). However, the start and end of the active season is usually not determined by temperature directly. Instead, diapause is most often triggered by photoperiodic cues, as photoperiod tends to be a much more reliable cue of long-term mean climatic conditions – and hence expected future conditions – than the current temperature alone (Tauber et al. 1986, Leather et al. 1995). We therefore expect photoperiodic responses to have evolved so that organisms enter diapause at the time of the year when the long-term mean temperature falls at, or slightly above, the lower developmental threshold.

Based on this, we used the months when the mean temperature falls above the lower developmental threshold temperature for insects (T_dmin_) as a biologically plausible estimate of the active period of a population. Doing so ensures that shorter periods of cold temperatures that occur within the active period will count toward the thermal conditions experienced by active insects, whereas warmer periods that occur outside these months will not (because an individual will then be in diapause).

In a review, including 66 species in 8 insect orders, Dixon et al. (Dixon et al. 2009) reported a mean T_dmin_ of 13.3 °C across all insects. Analyzing data from the original studies, we explored whether T_dmin_ changed systematically with latitude. We were able to retrieve data on the latitude of collection for 36 of the species in Dixon et al. (Dixon et al. 2009)(Supporting information, Table S1). A regression analysis of these 36 species showed no significant relationship between Tdmin and absolute latitude (r^2^ = 0.07, P = 0.12). However, there was a tendency for high latitude species to have somewhat lower T_dmin_ (slope = −0.08) than low latitude species. Mean and median T_dmin_ for species collected above or below 40°N in this data set was 8.9 and 11.4, respectively. Using the mean T_dmin_ from the original data in Dixon et al.(Dixon et al. 2009), i.e. 13.3 °C, could thus introduce a bias in that northern species would appear to be relatively more sensitive to warming. We therefore use the mid-range value between the mean and the median, 10°C, as a conservative estimate of T_dmin_ for all species when calculating the mean ambient temperature experienced during the active period. This estimate is approximately the same as that found in an earlier study by Buckley et al. (Buckley 2008).

### Habitat temperature during the active period (T_habA_)

Insects are ectotherms and therefore the body temperature of most species closely matches that of their habitat in the absence of behavioral thermoregulation (Angilletta 2009). Based on this, we calculated T_habA_ as the mean air temperature of those months were the temperature was above 10 °C for the 38 species of insects that were provided in Deutsch et al. (Deutsch et al. 2008). We were unable to retrieve the exact same observational climatic dataset as that used by Deutsch et al. (Deutsch et al. 2008). We therefore used a slightly different set of observational climatic data (the coordinates for the climate data set differ slightly at the collection location). To determine if our climatic dataset was comparable to that used by Deutsch et al. (Deutsch et al. 2008), we re-calculated T_hab_ using our climatic dataset and compared the results with those originally presented by Deutsch et al. (Deutsch et al. 2008). The results were highly congruent (Supporting information, Table S2).

### Climate data

Monthly baseline climate data for the 20^th^ century (1901-2009) for each month was obtained from (https://www.globalspecies.org/weather_stations/climatemap). For the future global warming we used the predicted mean monthly temperate for the year 2080 using, as in Deutsch et al (Deutsch et al. 2008), the simulation from the Geophysical Fluid Dynamics Laboratory model CM2.1 (Delworth et al. 2006) that is forced by the A2 greenhouse gas emissions scenario (obtained at http://www.climatewizard.org/).

### Warming tolerance (WT) and Thermal safety margin (TSM) for the active period

Following Deutsch et al. (Deutsch et al. 2008), thermal performance was estimated as the intrinsic rate of population growth (r_max_) across a range of constant temperatures. This value is a direct estimate of Darwinian fitness, and is thus a proper fitness estimate for each species (Stearns 1992). T_min_, T_opt_ and T_max_ for 38 species at the location of collection was obtained from the data set in Deutsch et al. (Deutsch et al. 2008). Thereafter, we estimated warming tolerance (WT) and thermal safety margin (TSM) at the site of collection for each species using the formulas WT = T_max_−T_hab_, and TSM = T_opt_−T_hab_ respectively, using the predicted temperature in 2080. As stated above, we calculated the yearly habitat temperature in two ways: using a 12-month approach considering each month mean (T_hab_; same as in Deutsch et al. (Deutsch et al. 2008)), and our new approach using only means of months where the temperature was >10 °C (T_habA_).

The effects of warming on insect performance was visualized by plotting WT and TSM against latitude of origin for each of the 38 species analyzed by Deutsch et al. (Deutsch et al. 2008) for their 2080 warming scenario. We created two different plots: one for the entire year (T_hab_) and one for only the active period (T_habA_). In these plots we also show the predicted temperatures increase at each latitude for the year 2080. To visualize trends in WT and TSM across latitude we used a second-degree polynomial model. To evaluate if tropical species were more sensitive to global warming compared to temperate species we calculated two temperature sensitivity indices: (1) WT - predicted mean temperature (ΔT in Figure 1); and (2) TSM – predicted mean temperature increase. Again, we did so using both T_hab_ and T_habA_. We define the tropics as the area between 23.5°N and 23.5°S (Osborne 2000). The differences in temperature sensitivity indexes between tropical and temperate species were tested with t-tests.

## Results

As originally reported by Deutsch et al. (Deutsch et al. 2008) and others, warming tolerances (WT) and thermal safety margins (TSM) increased steeply with latitude (Figure 2a,b) when we calculated these metrics using the annual mean habitat temperature T_hab_ as in Deutsch et al.’s (2008) original analysis. The pattern was significantly better described by a quadratic regression model than a linear model (WT: *F*_1,35_= 22.37, *P* < 0.001; TSM: *F*_1,35_= 29.19, *P* < 0.001). Similar to previous studies, tropical species (below 23° latitude) had on average significantly narrower warming tolerances (t-test, t= 3.62, *P*= 0.003) and thermal safety margins (t= 2.83, *P* = 0.015) than temperate species (above 23° latitude). Furthermore, one tropical species and one temperate species was predicted to experience mean annual temperatures that exceeded their thermal safety margin by 2080 (dots below the red line in Figure 2b). Because the data set is heavily biased towards temperate species, this represent 17% of tropical species and 1 % or temperate species being identified as at risk from warming.

In contrast, WT and TSM showed a considerably flatter latitudinal trend when we accounted for latitudinal differences in insect activity periods by calculating these metrics using the mean temperature during the active season T_habA_ (Figure 2c,d). The pattern was still significantly better described by a quadratic regression model than a linear model (WT: *F*_1,35_= 10.15, *P*= 0.003; TSM: *F*_1,35_= 11.21, *P*= 0.002). However, after accounting for activity periods, tropical and temperate species did not differ overall in either warming tolerance (Figure 3A; t-test, *t*= 1.30, d.f.=8.9, *P*= 0.22) or thermal safety margin (Figure 3B; t-test, *t*= 0.63, d.f.=7.43, *P*= 0.55) based on currently available data. Moreover, one tropical species and three temperate-zone species were predicted to experience mean temperatures during the active season that exceeded their thermal safety margins by 2080 (Figure 2D). This analysis accounting for latitudinal differences in insect active periods thus identified 17% of tropical species and 9% of temperate species as at risk to warming.

**Figure 3.**
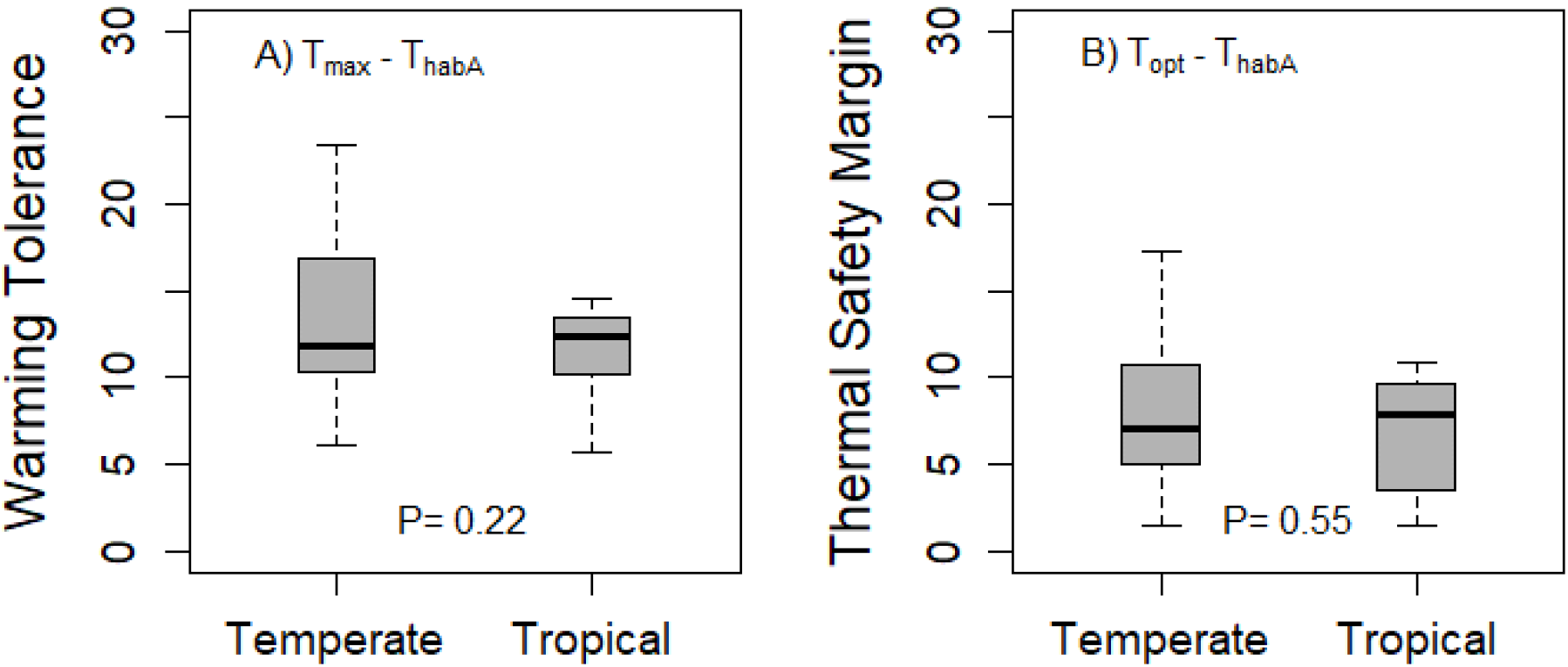
Difference in warming tolerance and thermal safety margins between temperate and tropical species when accounting for latitudinal differences in insect activity periods. Box plots show median, quartiles and range. P-values are based on t-tests. Tropical areas are defined as regions between 23°N and 23°S latitude.

Accounting for insect activity periods had a greater impact on the predicted warming tolerances and safety margins in temperate than tropical species. Warming tolerances calculated using the mean annual temperature T_hab_ were significantly higher than those calculated using the mean temperature during the active season T_habA_ for temperate (t-test, *t*= 3.00, d.f.=53.3, *P*= 0.004) but not tropical species (*t*= −0.008, d.f.=10, *P*= 0.99). Likewise, thermal safety margins were significantly higher when calculated using T_hab_ than T_habA_ for temperate (*t*= 3.22, d.f.=53.3, *P*= 0.002) but not tropical species (*t*= −0.007, d.f.=10, *P*= 0.99).

## Discussion

Our study suggests that temperate and tropical insects are more similar in their vulnerability to climate warming than expected based on several recent models. These findings challenge the view that temperate species are relatively safe from warming because they live in environments where the mean habitat temperature falls well below their current thermal optima. Specifically, we find that previous analyses that have compared thermal optima and maxima (T_opt_ and T_max_) with the annual mean habitat temperature (T_hab_) have greatly overestimated the warming tolerances (WT) and thermal safety margins (TSM) of temperate species (Figure 2A,B), but not those of tropical species. When we account for differences in seasonal activity patterns across latitude, by calculating the mean temperature during the months of the year when insects are developmentally active T_habA_, increases in mean WTs and TSMs with latitude becomes considerably smaller in magnitude than reported in several earlier studies (Figure 2C,D). This discrepancy simply arises because our metric T_habA_ is not biased downward by including very cold winter temperatures that are not experienced by developmentally active insects at higher latitudes.

Our analysis highlights the striking variability in warming tolerances and thermal safety margins in temperate species (Figure 2C,D). This variability, together with the weak latitudinal trend in WT and TSM, does not support the generalization that tropical species are more vulnerable overall than temperate species (Figure 3), although it should be noted that on average, TSM and WT tends to increase non-linearly with latitude here as in previous studies. The weak differences we observe between temperate and tropical taxa contrasts with the results from several earlier studies (Stillman and Somero 2000, Compton et al. 2007, Tewksbury et al. 2008, Deutsch et al. 2008, Huey et al. 2009, Dillon et al. 2010, Duarte et al. 2012) that have not accounted for activity periods.

We find the variation in TSM in the temperate zone to be so great that three temperate species – but only one tropical species – showed reduced fitness following the 5°C increase in average temperatures forecasted by 2080 under IPCC scenario A2 (Figure 2D; points falling under the red line). This represents 9% and 17% of temperate and tropical species studied, respectively. These relatively similar level of risk stands in stark contrast with the predictions from Deutsch et al.’s (Deutsch et al. 2008) original analysis. While their analysis also showed large levels of variation in WT and TSM for temperate species (c.f. Figure 2B), the mean WT and TSM was estimated to be so high for temperate species that all insects above 40°N were predicted to experience increased fitness even after a 10°C increase in temperature. Clearly, more data on both tropical and high-latitude species is needed before any firm conclusions can be drawn.

As a consequence of the variability observed among temperate species, our results suggest that global warming may have more complex ecological consequences in high-latitude environments. Thus, while warming may have a more uniformly negative impact on most species in the tropics, temperate areas may experience drastic increases in performance and fitness for some species, but drastic declines for others (Figure 2D). This could potentially have dramatic effects for the structure and functioning of these ecosystems (Tylianakis et al. 2008). Clearly, more focus should be directed toward identifying at-risk species in the temperate zone and forecasting the consequences to ecosystems from changes in the distribution and abundance of these species.

The effects of global warming will not only be felt through changes in mean temperature, but also from changes in the variance of temperature experienced over annual and diurnal time scales. In the analyses presented here, we focus on changes in mean temperature for two reasons. Firstly, it enables us to clearly isolate the magnitude of the effect of accounting for insect activity periods across latitude, and secondly, it enables us to directly compare the results from our analyses with those in the original, highly influential, study by Deutsch et al. (2008). However, our findings, that mid-latitude face a greater risk from warming than commonly expected, reinforce those of two recent studies that have re-analyzed the Deutsch et al. (2008) dataset to explore the consequences of increases in both the mean and variance of temperature under climate change (Kingsolver et al. 2013, Vasseur et al. 2014). Both these studies found that, because mid-latitude environments experience considerably greater variance in temperature than tropical environments, insects from mid-latitudes will experience greater decline in fitness under global warming because they will encounter temperatures above Topt with increasing frequency (Kingsolver et al. 2013, Vasseur et al. 2014). However, note that this arise because temperate environments show greater variation in in temperature than tropical environments. Our results add to these findings by showing that active insects also experience these temperate environments as *warmer* than expected based on conditions averaged over the whole year, because temperate insects have shorter active periods. When we add the effect of thermal fluctuations (and increases in thermal variability under warming) to this, we expect that this will further exacerbate the vulnerability of the temperate species that we identify as having narrow thermal safety margins.

The study by Kingsolver et al. (2013) is an especially relevant comparison, as they also performed separate analyses using temperature data for both the full year and the ‘growing season’, which they define as May-October in the northern hemisphere (but note that their analysis did not account for differences in the length and timing of the active period across latitude, as we do here). As in our study, Kingsolver et al. (2013) found that a considerably greater proportion of mid-latitude species (20-40° absolute latitude) will experience declines in fitness when we only consider the conditions they experience during the growing season. In addition, Kingsolver et al. (2013) also found that responses were especially heterogeneous at mid-latitudes, with some species showing increases, and some species decreases, in fitness, which is in concert with our findings.

Estimates of WT and TSM from TPCs for fitness (*r*) represent a highly valuable source of data on species’ vulnerabilities to warming as they have been estimated for a relatively large number of species using a consistent methodology, and capture the overall effect of temperature on a set of complex underlying processes that are intrinsically linked to the demographic rates of populations. However, it is becoming increasingly clear that more complex models will be needed to predict the response of individual species with any certainty. Mechanistic models, tailor-made for specific organisms, are becoming increasingly sophisticated. These incorporate, for example, the effects of thermoregulatory behavior (Buckley et al. 2015), capacity for thermal acclimation (Nilsson-Örtman and Johansson 2017), response to the duration and intensity of extreme temperatures (Marshall and Sinclair 2015), genetic (co)variances (van Heerwaarden et al. 2016) and interspecific interactions (Nilsson-Örtman et al. 2014, Diamond et al. 2016). However, to apply such mechanistic models on a global scale will require a large amount of data that is currently not available for more than a handful of species. Therefore, we believe that TPCs for fitness will continue to serve an important role for identifying the broad trends in the vulnerability to warming. Going forward, we strongly urge researchers to consider latitudinal differences in active periods when exploring this rich source of information.

Our analysis ignores sources of selection that occurs during the non-active period. This decision may be defensible when analyzing changes in the expression of performance traits (feeding, growth, locomotion, etc) that are only expressed during this part of the year. However, because fitness represents the joint outcome of survival and reproductive success, it may be especially important to also account for mortality that occur during the non-active periods. Likewise, because TPCs for fitness are typically estimated under controlled laboratory conditions, it is important to note that published fitness estimates also ignore many other important sources of selection, including predation, competition, resource availability, etc., that occur both within and outside the active period.

Finally, we note that the information currently available on insect thermal responses is highly incomplete and biased. Large areas remain entirely unsampled (Supporting information, Figure S4), most notably all of South America, and very few populations above 50°N have been studied (none above 53°N). Furthermore, the taxonomic and ecological characteristics of the studied species remain highly limited. Specifically, of the 38 species for which TPCs for fitness have been estimated, 24 are classified as pest species and 10 are used for biological control (Supporting information, Table S3). Of the four species remaining, one is a springtail that is dominant in rice fields and three are aphids feeding on ornamental trees and thistles. It seems highly probable that this suite of species shares several life history characteristics – and possibly thermal adaptations – as a consequence of having successfully exploited human crops (and our attempts to defend against them) that may make them poor representatives of all insects.

In summary, our work reveals that temperate species have considerably narrower safety margin to warming than suggested by several earlier analysis. Because temperate ectotherm species also harbor considerable variation in T_opt_ and T_max_ (Supporting information, Table S2) it will be important to have much more data on thermal tolerance and fitness components across latitudes before we can make more precise predictions on the impacts of global warming.

## Acknowledgements

We thank Lock Rowe for valuable comments on a previous version of this article.

## Author contributions

FJ conceived the study and wrote the first draft. All authors contributed equally to data collection data analysis and writing.

## Permit(s)

No permits were needed.

Supplementary information is available for this paper at https://doi.org/xxx

