## Supplementary material S1 for "Impact of global warming on insects: are tropical species more vulnerable than temperate species?"

Supporting information. Table S1.

$T_{dmin}$ for the insects used to calculate average $T_{dmin}$ and regression of $T_{dmin}$ against latitude. Note that more species are available in Dixon *et al.* (2009: *Functional Ecology* 23: 257-264), but we were not able to retrieve the origin of latitude for those species.

Lowest development temperature in insects
Data from Dixon *et al.* 2009. *Functional Ecology* 23: 257-264

| **Insect order** | **Species** | **Latitude North** | **Temperature** |
|---|---|---|---|
| Thysanoptera | *Ceratothripoides claratris* | 13 | 16,3 |
| Psocoptera | *Liposcelis bostrychophila* | 29 | 10,7 |
| Neuroptera | *Anomalochrysa frater* | 21 | 7,8 |
| Lepidoptera | *Sesamia nonagrioides* | 42 | 11,9 |
| Lepidoptera | *Plutella xylostella* | 30 | 9,9 |
| Lepidoptera | *Maruca vitrata* | 7 | 10,9 |
| Lepidoptera | *Lacanobia subjuncta* | 47 | 6,7 |
| Lepidoptera | *Endopiza viteana* | 42 | 9,0 |
| Lepidoptera | *Carposina sasakii* | 37 | 10,5 |
| Hymenoptera | *Venturia canescens* | 38 | 12,0 |
| Hymenoptera | *Muscidifurax raptorellus* | 40 | 12,3 |
| Hymenoptera | *Muscidifurax raptor* | 44 | 11,4 |
| Hymenoptera | *Muscidifurax zaraptor* | 50 | 13,3 |
| Hymenoptera | *Cirrospilus* | 39 | 6,0 |
| Hymenoptera | *Oomyzus sokolowski* | 30 | 12,0 |
| Hymenoptera | *Aphidius gifuensis* | 35 | 5,5 |
| Hemiptera | *Pemphigus populitransversus* | 26 | 6,0 |
| Hemiptera | *Myzus persicae* | 30 | 4,8 |
| Hemiptera | *Metopolophium dirhodum* | 52 | 2,0 |
| Hemiptera | *Macrosiphum avenae* | 52 | 1,7 |
| Hemiptera | *Macrolophus pygmaeus* | 38 | 8,9 |
| Hemiptera | *Lipaphis erysimi* | 30 | 6,8 |
| Hemiptera | *Bemisia tabaci* | 36 | 9,7 |
| Hemiptera | *Aphis spiraecola* | 26 | 1,7 |
| Hemiptera | *Acyrthosiphon pisum* | 54 | 4,8 |
| Hemiptera | *Abgrallaspis cyanophylli* | 51 | 12,3 |
| Diptera | *Stomoxys calcitrans* | 50 | 12,0 |
| Diptera | *Lydella jalisco* | 20 | 14,0 |
| Diptera | *Culiseta melanura* | 39 | 10,0 |
| Diptera | *Chironomus crassicaudatus* | 28 | 11,3 |
| Coleoptera | *Pterostichus nigrita* | 51 | 6,7 |
| Coleoptera | *Euhrychiopsis lecontei* | 45 | 11,7 |
| Coleoptera | *Prostephanus truncatus* | 19 | 12,3 |
| Coleoptera | *Nephus includens* | 39 | 11,0 |
| Coleoptera | *Nephus bisignatus* | 39 | 9,8 |
| Coleoptera | *Agasicles hygrophila* | 36 | 12,8 |
