## Supplementary material S3 for "Impact of global warming on insects: are tropical species more vulnerable than temperate species?"

Supporting information. Table S3.

Geographical position, systematic position, and ecological characteristics of the 38 species for which TPCs for fitness have been estimated. For a list of full references see Deutsch *et al.* 2008. *Proc. Natl. Acad. Sci.* 105: 6668–6672.

| Lat | Long | Order | Species | Pest | Ecology | Reference |
|---|---|---|---|---|---|---|
| 46.98 | -71.48 | Coleoptera: Coccinellidae | Stethorus punctillum | Biological control | Predator on spider mites | Roy et al 2003 |
| -31.08 | 150.93 | Hemiptera: Coccoidea: Dactylopiidae | Dactylopius austrinus | Biological control | Feeds on invasive cacti | Hosking 1984 |
| 38 | 23 | Hemiptera: Miridae | Macrolophus pygmaeus | Biological control | Generalist predator used to control greenhouse pests | Perdikis and Lykouressis 2002 |
| 25 | 68 | Hymenoptera: Braconidae | Cotesia flavipes | Biological control | Parasitizes a crambid moth pest of cereal crops | Mbapila and Overholt 2001 |
| 30.25 | 120.2 | Hymenoptera: Braconidae | Cotesia plutellae | Biological control | Parasitizes on a moth pest on cruciferous crops | Shu-sheng et al 2000 |
| -3 | 40 | Hymenoptera: Braconidae | Cotesia sesamiae | Biological control | Parasitizes on cereal stem borers | Mbapila and Overholt 2001 |
| 45 | -93.17 | Hymenoptera: Pteromalidae | Muscidifurax raptor | Biological control | Parasitizes on muscoid flies incl. Stomoxys calcitrans | Lysyk 2000 |
| 40.82 | -96.67 | Hymenoptera: Pteromalidae | Muscidifurax raptorellus | Biological control | Parasitizes on muscoid flies incl. Stomoxys calcitrans | Lysyk 2001 |
| 49.9 | -112.9 | Hymenoptera: Pteromalidae | Muscidifurax zaraptor | Biological control | Parasitizes on muscoid flies | Lysyk 2001 |
| 13.48 | 2.17 | Hymenoptera: Trichogrammatidae | Uscana lariophaga | Biological control | Parasitizes on a pest on cowpeas | Huis et al 1994 |
| -26.5 | 31.5 | Coleoptera: Chrysomelidae: Bruchinae | Callosobruchus rhodesianus | Pest | Feeds on stored legumes | Giga and Smith 1983 |
| -26 | 150 | Coleoptera: Tenebrionidae | Pterohalaeus alternatus | Pest | Soil-dwelling larvae damage crops | Allsopp 1981 |
| -27.47 | 151.48 | Coleoptera: Tenebrionidae | Pterohelaeus darlingensis | Pest | Soil-dwelling larvae damage crops | Allsopp 1981 |
| 49.68 | -112.8 | Diptera: Muscidae | Stomoxys calcitrans | Pest | Adult flies suck blood from mammals | Lysyk 1998 |
| 23.12 | 113.25 | Diptera: Tephritidae | Bactrocera dorsalis a | Pest | Feeds on fruits and vegetables | Yang et al. 1994 |
| 20.88 | -156.52 | Diptera: Tephritidae | Ceratitis capitata | Pest | Feeds on fruits | Vargas et al 1997 |
| 25.47 | -80.48 | Hemiptera: Aleyrodidae | Bemisia argentifolia | Pest | Feeds on a range of crops | Wang and Tsai 1996 |
| 51.53 | 0 | Hemiptera: Aphididae | Acyrthosiphon pisum | Pest | Feeds on legumes | Morgan et al 2001 |
| 34.98 | 138.4 | Hemiptera: Aphididae | Aphis citricola | Pest | Feeds on fruit trees | Komazaki 1982 |
| 36.07 | 114.22 | Hemiptera: Aphididae | Aphis gossypii | Pest | Feeds on cotton | Xia et al 1999 |
| 38.93 | -92.3 | Hemiptera: Aphididae | Brevicoryne brassicae | Pest | Feeds on Brassicaceae crops | DeLoach 1974 |
| -30.51 | 151.66 | Hemiptera: Aphididae | Eriosoma lanigerum | Pest | Feeds on apple trees | Asante et al 1991 |
| 38.93 | -92.3 | Hemiptera: Aphididae | Hyadaphis pseudobrassicae | Pest | Feeds on Brassicaceae crops | DeLoach 1974 |
| 38.93 | -92.3 | Hemiptera: Aphididae | Myzus persicae | Pest | Feeds on peach trees | DeLoach 1974 |
| 26.14 | -81.8 | Hemiptera: Aphididae | Rhopalosiphum rufiabdominal | Pest | Feeds on rice, bananas, etc. | Tsai and Liu 1998 |
| -42 | 147 | Hemiptera: Aphididae | Sitobion fragariae | Pest | Feeds on leaves of blackberries (Rubus sp.), etc. | Turak et al 1998 |
| -33 | 152 | Hemiptera: Aphididae | Sitobion miscanthi | Pest | Feeds on cereals | Turak et al 1998 |
| 34.98 | 138.4 | Hemiptera: Aphididae | Toxoptera citricidus aurantium | Pest | Feeds on citrus | Komazaki 1982 |
| -21.09 | 149.09 | Hemiptera: Coccoidea: Pseudococcidae | Saccharicoccus sacchari | Pest | Feeds on sugarcane | Rae and De'ath 1991 |
| 6.45 | 2.35 | Hemiptera: Heteroptera: Coreidae | Clavigralla shadabi | Pest | Feeds on pods of cowpea | Dreyer and Baumgartner 1996 |
| 26.2 | -80.1 | Hemiptera: Psylloidea: Liviidae | Diaphorina citri | Pest | Feeds on citrus | Liu and Tsai 2000 |
| 26 | -98.8 | Lepidoptera: Crambidae | Diatraea lineolata | Pest | Feeds on Maize | Rodriguez-del-Bosque et al 1992 |
| 33.4 | -117.2 | Thysanoptera: Thripidae | Scirtothrips perseae | Pest | Feeds on Avocado | Hoddle 2002 |
| 35.36 | 132.75 | Thysanoptera: Thripidae | Thrips tabaci | Pest | Feeds on onions and garlics | Murae 2000 |
| 37 | 127.5 | Collembola: Onychiuridae | Paronychiurus kimi | | Dominant springtail in rice fields | Choi et al 2002 |
| 52.4 | 1.18 | Hemiptera: Aphididae | Drepanosiphum acerinum | | Feeds on leafs of sycamore trees | Wellings 1981 |
| 52.4 | 1.18 | Hemiptera: Aphididae | Drepanosiphum platanoids | | Feeds on leafs of sycamore trees | Wellings 1981 |
| -35.3 | 149.08 | Hemiptera: Aphididae | Hyperomyzus lactucae | | Feeds on sowthistle (Asteraceae: Sonchus) | Shu-sheng and Hughes 1987 |
