## Supplementary material S4 for "Impact of global warming on insects: are tropical species more vulnerable than temperate species?"

Supporting information. Figure S4

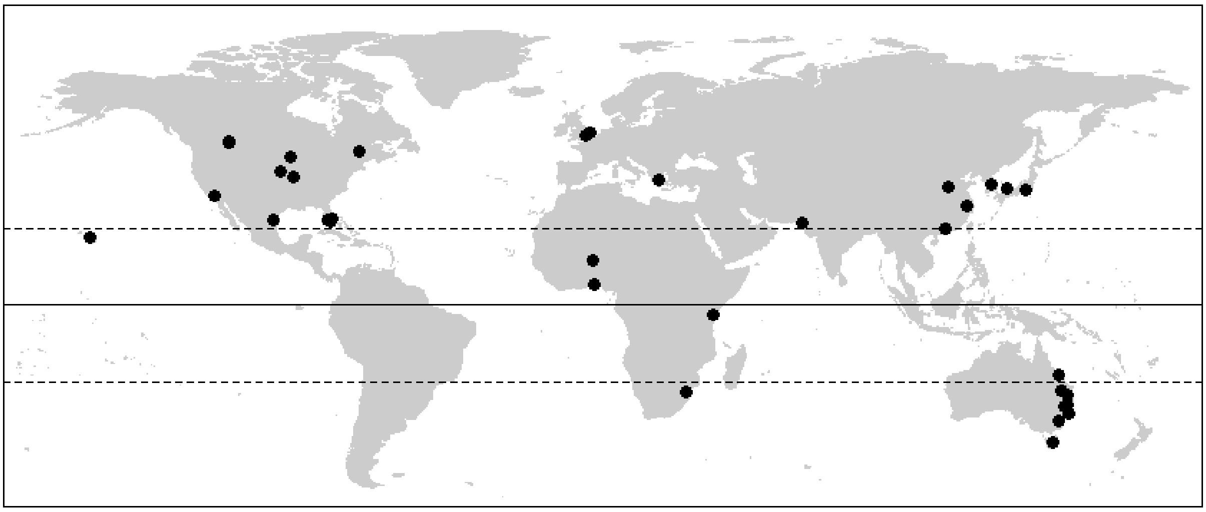

**Figure S4.** Geographic location of the origin of the 38 insect species for which thermal performance curves (TPCs) for fitness have been estimated in this study.
